# Transcriptome-based Analysis of Tomato Genotypes Resistant to Bacterial spot (*Xanthomonas perforans)* Race T4

**DOI:** 10.1101/736660

**Authors:** Rui Shi, Dilip R. Panthee

**Affiliations:** Department of Horticultural Science, North Carolina State University, Mountain Horticultural Crops Research & Extension Center, 455 Research Drive, Mills River, NC 28759, USA; Department of Crop and Soil Science, North Carolina State University, Raleigh, NC 27695-7620, USA

**Keywords:** RNAseq, tomato, Bacerial spot, Xanthomonas perforans Race T4, inoculation, resistance, differential expressed gene, SNP/INDELs, chromosome, QTL

## Abstract

Bacterial spot (BS) is one of the most devastating foliar bacterial diseases of tomato caused by multiple species of *Xanthomonas*. We performed the RNA-Seq analysis of three tomato lines with different level of resistance to *Xanthomonas perforans* race T4 to study the differentially expressed genes (DEGs) and transcript-based sequence variations.

Analysis between inoculated and control samples revealed that resistant line PI 270443 had more DEGs (834), followed by susceptible line NC 714 (373), and intermediate line NC 1CELBR (154). Gene functional analysis based on *Gene Ontology* (GO) terms revealed that more GO terms (51) were enriched for up-regulated DEGs in the resistant line PI 270443, and more down-regulated DEGs (67) were enriched in the susceptible line NC 714. The specific analysis for DEGs in biotic stress pathway using MapMan software showed more up-regulated biotic stress pathway DEGs (67) for PI 270443 compared to more down-regulated DEGs (125) for susceptible NC 714 line. One interesting feature was that resistant PI 270443 has three up-regulated DEGs for PR-protein, and susceptible line NC 714 has one down-regulated R gene, which is disease-related.

Analysis of sequence variations called from RNA-Seq reads against the reference genome of susceptible Heinz 1706 showed that chr11 which has multiple reported resistance QTLs to BS race T4 is identical between two resistant lines, PI 270443 and NC 1CELBR, suggesting that these two lines share the same resistance QTLs on this chromosome. Several loci for PR-resistance proteins with sequence variation between the resistant and susceptible tomato lines were identified near the known *Rx4* resistance gene on chr11. These findings may be useful for further molecular breeding of tomato.

## Introduction

Bacterial spot (BS) is a foliar disease of tomato caused by multiple species of *Xanthomonas*. There are four physiological races of *Xanthomonas*, including races T1 to T4, which are distributed throughout the world, particularly in warm and humid regions and causing a significant yield loss every year. Association between race and species is classified as *X. euvesicatoria* (race T1; *Xe*), *X. vesicatoria* (race T2; *Xv*), *X. perforens* (race T3 and T4; *Xp*), and *X. gardneri* (no race designations described so far; *Xg*) [1, 2].

Resistance to BS is both monogenic and polygenic conferring partial resistance [2]. Multiple resistance genes and loci for BS have been identified in tomato and has been summarized by Pei, Wang [3]. ‘Hawaii 7998’ is a differential line for identifying race T1, which are *Xanthomonas* spp — carrying the *AvrRxv* gene [4]. ‘Hawaii 7998’ remains the most reliable source of resistance to race T1 [3, 5], which is conferred by three independent loci (*rx1* and *rx2* on opposite arms of chromosome 1 and *rx3* on chromosome 5) and may be modified by three susceptible loci on chromosomes 3, 9 and 11 [6]. The dominant allele *Rx3* confers resistance in the field, explaining 41% of the phenotypic variation [7] and is considered to be the most effective loci for T1 resistance. It is not clear yet whether *rx3* and *Rx3* are alleles of the same gene, or closely associated genes on the same chromosome [3].

Resistance to race T2 has been documented in ‘Hawaii 7983’ that expressed partial resistance over multiple seasons [8]. As *Xp* contains the major races (T3 and T4), which cause many annual epidemics in tomato growing regions [2, 9], much work has been done to identify and breed resistance against this group of the *Xanthomonas*.

Race T3 resistance has been identified in several lines including ‘Hawaii 7981’, and *S. pimpinellifolium* accessions PI 126932 and PI 128216, each conferring a HypR in the presence of the pathogenic expression of the *avrXv3* gene, and partial resistance in field assessments [10]. T4 resistance has been identified in LA 716 (*S. pennellii*), being conferred by *Xv4*. The previously mentioned line PI 114490 also harbors strong resistance to race T4.

Mapping and characterization of loci help to develop resistant breeding lines. QTL associated with BS race T4 were identified in the populations derived from Hawaii 7998 and PI 114490 lines and mapped to chromosome 11 and chromosome 3, respectively [11].

Resistance to Bacterial spot is very complex, and made the introgression of resistance into the desirable genetic background very challenging. Although the QTLs with the reasonably high level of *R^2^*-value (29.4% and 4.8%, respectively from chromosomes 11 and 3) to BS race T4 had been identified using QTL mapping method [11], its introgression into the breeding lines to achieve the desired level of BS resistance has not been achieved. In this context, it is reasonable to investigate the gene regulation network in more details, so that the resistance mechanism can be understood better. Differential gene expression analysis by conducting RNA-Seq experiments has been used to understand such mechanism in different species [12]. Recently, Du, Wang (13] investigated the expression profiles of genes in response to BS race T3 infection, and found that 78 genes were up-regulated in PI 114490 (resistant parent) and 15 genes were up-regulated in OH 88119 (susceptible parent) six days post-inoculation (dpi). With that information available, it is logical to investigate the genes specially expressed on exposure to BS race T4 in tomato, which has not been reported to date. With the availability of detailed annotation for most of the genes in the tomato genome, functional classification of the genes and pathway analysis has been much more comfortable now. Gene expression analysis approach using RNA-Seq analysis technology has been applied to unravel the gene function and defense mechanism in tomato, soybean and several other plant species [13, 14]. This approach is useful to identify the gene(s) associated with complex traits by comparing the detailed network of gene regulation between host and pathogens and eventually determining the phenotypic trait [12, 15–17]. Hence this approach is suitable to identify the gene network involved in conferring resistance to the BS.

Available tomato genome sequence [18] also presents opportunity to identify sequence polymorphism in the form of Single Nucleotide Polymorphisms, Insert and Deletion (SNP/INDELs) genome-wide, which can be developed into molecular markers for breeding program [19–21]. RNA-Seq data is a valuable resource for the detection of SNP/INDELs from gene transcripts, a sub-set of the whole genome sequence, but can be more related to functional analysis. For instance, to detect non-synonymous SNP/INDELs, which lead to gene function alternation due to amino acid sequence change [20].

In North Carolina, *X. perforans* race T4 is the predominant race [22]. There are limited QTL analysis information and genetic dissection information available as of now. In this situation, we evaluated a few tomato breeding lines for BS resistance. Based on this screening, we selected three lines with a relatively good level of resistance and susceptible to BS. These lines were used for transcriptome-based analysis in this study.

## Material and methods

### Plant materials

Thirty tomato lines grown in the greenhouse were evaluated for their resistance to bacterial spot (*X. perforans*). These tomato lines were sown in 4P soil mixture (Fafard®, Florida, USA) in 24-cell trays in the greenhouse at the Mountain Horticultural Crops Research & Extension Center, Mills River, NC. Six plants per genotype were planted in two replications in a completely randomized design. Plants were fertilized using a mixture of fertilizer containing a ratio of 20:20:20 of nitrogen, phosphorus, and potassium, respectively. Standard greenhouse treatments for insects and fungal diseases management were used, but copper was not applied to control the bacterial diseases.

Based on the disease score, three tomato lines PI 270443, NC 1CELBR, and NC 714 were selected to be used for the RNA-Seq analysis. PI 270443 (*Solanum pimpinellifolium* L.), a small-fruited line, was found to have the least level of BS and used as a resistant line in this study. NC 714 is a large-fruited tomato breeding line with excellent horticultural traits developed from NC State breeding program [23]. It developed the most BS disease and was used as a susceptible line in this study. NC 1CELBR is also a large-fruited late blight resistant tomato breeding line developed from NC State University tomato breeding program [24]. Late blight resistance comes from one of the *S. pimpinellifolium* (LA3707) line. It had a medium level of BS disease resistance and was used as a tomato line in this study.

### Bacterial spot inoculation and disease evaluation

Plants were artificially inoculated with Isolate 9 of *X. perforans*, which was found to be extremely virulent. This is a field isolate collected from infected tissue of tomato plant in western NC and characterized as *X. perforans* race T4 using differential tomato lines [5, 25–27] by Dr. Jefferey B. Jones lab, University of Florida, Gainesville, Florida. The strain was maintained in pure culture and stored at −80°C. The isolate was grown in Yeast Dextrose Chalk (YDC) agar medium [28] at 28°C for 24-48 hours and was then overlaid with sterile distilled water. The bacteria were dislodged from the plates and the resulting bacterial suspensions were pooled in a sterile glass container. The suspension was standardized by determining its optical density at 600 nm using an LKB Biochrom Ultrospec II Spectrophotometer (American Laboratory Training, USA), and diluted as needed to obtain an OD_600_ of 0.3 (approximately 2-5×10^8^ CFU/mL). Diluted cells were immediately used for inoculations.

For greenhouse inoculations, humidity around the plants was maintained using V5100NS humidifiers (Vicks Ultrasonic Humidifiers, Hudson, NY, USA) from 24 hours before inoculation to 48 hours after inoculation and by covering the seedlings with clear plastic. Four to six weeks after transplanting, the seedlings were sprayed with the bacterial suspension till foliar runoff using a hand sprayer around sunset. Sterile water was used for mock inoculation. Leaf tissue samples after inoculation were collected in liquid nitrogen and stored at −80°C until further processed.

Greenhouse plants were scored for foliar symptoms on the most severely infected leaves using Horsfall-Barratt scale where 0% = 1, 1-3% = 2, 3-6% = 3, 6-12% = 4, 12-25% = 5, 25-50% = 6, 50-75% = 7, 75-87% = 8, 87-94% = 9, 94-97% = 10, 97-100% = 11 and 100% dead tissue = 12.

### RNA extraction and RNA-Seq library construction

Leaves from three breeding lines were collected in liquid nitrogen and three replicates at 48 hours after inoculation (hai). Frozen samples were stored at −80°C freezer. Before RNA extraction, frozen leaves were ground into fine powder by using pestle and mortar in liquid nitrogen. About 100 mg ground tissue sample was transferred into 1.5 ml tube for the extraction of total RNA using the Qiagen Plant RNeasy mini kit (Qiagen, Hilden, Germany). The RNA quality and quantity were evaluated by using Nanodrop (Fisher Scientific, Waltham, MA) and MOPs gel electrophoresis [29]. Total RNA samples were used for RNA-Seq library construction using NEBNext^®^ Ultra^TM^ Directional RNA Library Prep Kit for Illumina (New England BioLabs, Ipswich, MA, USA). We followed the protocol for normal insertion size option.

### RNA-Seq deep sequencing, data processing, mapping and differential gene expression analysis

RNA-Seq data was generated on Illumina HiSeq2500 instruments in 150 bp read mode at Genomic Sciences Laboratory, North Carolina State University, Raleigh, NC and reads were provided in FASTq format. All reads were quality checked using FastQC [30], and trimmed to eliminate poor quality bases (Q30) using fastq-mcf function.

Reads were mapped against tomato Heinz 1706 genome assembly SL3.0 using Hisat2 version 2.1.0 [31]. Reads mapped to gene assembly were manipulated using Samtools [32] for sorting/indexing, and raw count of reads mapped to annotated gene model (ITAG3.2 version) was extracted using Bedtools version 2.25.0 [33]. Information of raw counts of mapped RNA-Seq reads to annotated gene models were analyzed using EdgeR R package [34], including exploration of RNA-Seq libraries relationship using the plotMDS function, gene expression normalization followed TMM algorithm [35], and differentially expressed genes (DEG) identification followed classical approach. Criteria of DEG is gene expression fold change >2.0 times between compared group of samples, and statistics level in the form of false discovery rate (FDR) <0.05.

### Gene functional and pathway analysis

Biological function evaluation for DEGs was conducted using online GO analysis toolkit AgriGO2.0 [36] following an option of using ITAG3.2 version transcript ID and suggested ITAG3.2 background.

MapMan software (version 3.6.0RC1) was used to evaluate the DEGs’ function in the different pathway [37] using a list of DEGs with fold change in log2 between control and inoculated samples for each tomato line as input data. Gene annotation in the pathway analysis was prepared via Mercator online software within PlabiPD website (http://www.plabipd.de) based on ITAG3.2 protein sequence and followed the default annotation parameter, such as include database of TAIR Release 10, SwissProt/UniProt Plant Proteins, Clusters of orthologous eukaryotic genes database (KOG), conserved domain database, and set BLAST_CUTOFF as 80.

### SNP/INDEL identification

Sequence variations were extracted using the mpileup function of Samtools package [32] from mapping result file in the format as BAM. For even comparison, these BAM files were the result of mapping the same amount of clean RNA-Seq reads (20 million) from two control RNA-Seq libraries for each tomato line. To get high-quality SNP/INDELs, data in raw Variant Calling Format (VCF, version 4.0) files were filtered for minimum depth (DP) 10 and SNP/INDELs quality (Q) over 30 [20]. To screen SNP/INDELs specific to individual tomato line, loci with homozygous SNP/INDEL against Heinz 1706, i.e., in the form of 1/1 in genotype (GT) section of VCF output file), were selected.

Genes with SNP/INDELs were screened using vlookup function in Excel environment, and location information of genes with SNP/INDELs on the chromosome was converted into CSV format and used as input for R QTL package [38] to generate a genetic map for visualization. Online software Venny2.1 online [39] and BioVenn [40] were used to generate Venn diagrams or extract overlapping information for DEGs, SNP/INDELs.

## Results

### Three tomato lines with different level of resistance to BS were selected for RNA-Seq analysis

The resistance level of 30 tomato breeding lines was evaluated by inoculation with BS (*X. perforans*) race T4 in the greenhouse at Mountain Horticultural Crops Research and Extension Center, Mills River, NC. Leaf samples of these lines were collected and frozen in liquid nitrogen 48 hours after BS inoculation and then stored at −80°C before RNA extraction and RNA-Seq analysis.

The resistance level of these 30 tomato lines as shown in the calculated Area Under Disease Progress Curve (AUDPC) based on the data from greenhouse experiments was used as a single indicator to select the genotype for disease resistance. Among these tomato lines, PI 270443 was the most resistant line followed by PI 114490, CLN-2413A, LA 2093, LA4277, and Fla. 7060_Xv4 with an AUDPC value of less than 280 (Table 1). There was no significant difference between these lines for the level of bacterial spot resistance. Average disease incidence took place about 5.5 days after inoculation (dai) in LA2093, whereas it was 6 dai in PI 114490, LA4277, and Fla.7060_Xv4. In the case of PI 270443 and CLN2413A, the average disease incidence was 7 dai. Among the tomato lines, Heinz 1706 was the most susceptible line followed by NC 714, NC 6 Grape, NC EBR7, and Hawaii 7981 with an AUDPC value of more than 608 (Table 1). The time for the first disease symptom appearance was not much different in the group of susceptible lines, which ranged from 5.5 to 6 days. Even in the wide array of lines, NC 1CELBR was intermediate in its response to the BS along with other lines including NC 25P, NCEBR8, CLN-2418A, NC 161L, NC EBR6, Money Maker, LA2653, and NC123S. The level of disease development reported in terms of AUDPC in these lines ranged from 420 to 540 (Table 1). It seems that the disease develops quickly in the susceptible lines, whereas the rate of disease development is very slow in the resistant lines. Hawaii 7998, which is widely used as a resistant line to BS, was very close to NC 1CELBR in AUDPC value, whereas NC 714 was the most susceptible line.

**Table 1:** Bacterial spot disease development in the tomato breeding lines

| Genotype | First symptom dai | 6 dai (21DAP) | 7 dai (24DAP) | 9 dai (26DAP) | 11 dai (28DAP) | 13 dai (30DAP) | 15 dai (32DAP) | 17 dai (34DAP) | AUDPC |
| --- | --- | --- | --- | --- | --- | --- | --- | --- | --- |
| **PI 270443** | **7.0** | **1.0** | **1.0** | **4.5** | **10.0** | **22.5** | **20.0** | **17.5** | **135.5** |
| PI 114490 | 6.0 | 1.0 | 1.0 | 12.5 | 20.0 | 22.5 | 35.0 | 37.5 | 221.5 |
| CLN-2413A | 7.0 | 1.0 | 3.0 | 8.0 | 22.5 | 27.5 | 32.5 | 32.5 | 222.5 |
| LA2093 | 5.5 | 1.0 | 1.0 | 4.5 | 20.0 | 32.5 | 35.0 | 35.0 | 223.0 |
| LA4277 | 6.0 | 1.0 | 1.5 | 10.0 | 17.5 | 42.5 | 42.5 | 37.5 | 267.8 |
| Fla 7060_Xv4 | 6.0 | 1.0 | 3.0 | 10.0 | 15.0 | 35.0 | 50.0 | 50.0 | 279.0 |
| NC 25P | 6.0 | 1.0 | 3.0 | 17.5 | 32.5 | 37.5 | 45.0 | 47.5 | 321.5 |
| NCEBR8 | 6.5 | 1.0 | 3.0 | 17.5 | 25.0 | 45.0 | 50.0 | 42.5 | 326.5 |
| CLN-2418A | 6.0 | 1.0 | 10.5 | 20.5 | 35.0 | 40.0 | 45.0 | 42.5 | 351.3 |
| NC161L | 6.5 | 2.0 | 8.0 | 25.5 | 30.0 | 40.0 | 47.5 | 47.5 | 356.5 |
| NCEBR6 | 7.0 | 1.0 | 3.0 | 12.5 | 42.5 | 45.0 | 50.0 | 50.0 | 359.0 |
| Money maker | 6.0 | 1.0 | 9.5 | 27.5 | 35.0 | 45.0 | 52.5 | 52.5 | 397.8 |
| LA2653 | 5.5 | 1.5 | 10.0 | 17.5 | 50.0 | 45.0 | 50.0 | 47.5 | 399.8 |
| NC123S | 6.5 | 1.0 | 15.5 | 22.5 | 32.5 | 52.5 | 47.5 | 50.0 | 400.3 |
| VF36 | 6.5 | 10.0 | 5.5 | 25.5 | 35.0 | 47.5 | 52.5 | 52.5 | 402.3 |
| Pto-S | 5.5 | 1.0 | 6.0 | 17.5 | 40.0 | 60.0 | 57.5 | 55.0 | 421.5 |
| Hawaii 7998 | 6.0 | 1.5 | 15.5 | 30.5 | 45.0 | 47.5 | 50.0 | 35.0 | 422.0 |
| CLN-1466EA | 6.0 | 1.0 | 20.5 | 32.0 | 42.5 | 42.5 | 47.5 | 45.0 | 426.8 |
| **NC 1CELBR** | **6.0** | **1.5** | **16.0** | **32.5** | **47.5** | **37.5** | **50.0** | **55.0** | **432.3** |
| NC 30P | 7.0 | 1.0 | 10.5 | 20.5 | 40.0 | 57.5 | 57.5 | 55.0 | 433.8 |
| NC 1CS | 6.5 | 1.0 | 10.5 | 21.0 | 42.5 | 55.0 | 57.5 | 55.0 | 434.8 |
| NC 22L-1(2008) | 6.5 | 5.0 | 10.5 | 25.0 | 52.5 | 57.5 | 60.0 | 55.0 | 478.8 |
| HI7997 | 6.0 | 1.0 | 11.0 | 22.5 | 50.0 | 60.0 | 60.0 | 67.5 | 481.5 |
| Pto-R | 6.0 | 8.0 | 20.0 | 41.0 | 47.5 | 57.5 | 60.0 | 65.0 | 539.0 |
| Hawaii 7981 | 6.0 | 5.5 | 17.0 | 50.0 | 60.0 | 70.0 | 65.0 | 67.5 | 608.3 |
| NCEBR7 | 6.0 | 1.0 | 9.5 | 25.0 | 50.0 | 85.0 | 90.0 | 95.0 | 620.3 |
| NC 6Grape | 6.0 | 15.0 | 20.5 | 40.0 | 65.0 | 77.5 | 77.5 | 75.0 | 668.8 |
| **NC 714** | **5.5** | **6.0** | **15.0** | **40.0** | **65.0** | **85.0** | **87.5** | **85.0** | **686.5** |
| NC 5Grape | 5.5 | 8.0 | 25.0 | 55.0 | 67.5 | 72.5 | 80.0 | 80.0 | 704.5 |
| Heinz 1706 | 6.0 | 3.5 | 25.0 | 55.0 | 67.5 | 87.5 | 92.5 | 90.0 | 762.8 |

Based on the results presented in Table 1, three tomato lines were selected to represent a different level of resistance to BS for RNA extraction and subsequent RNA-Seq analysis. Among these lines, PI 270443 was the most resistant line, whereas NC 714 was one of the most susceptible tomato breeding lines, which has similar resistance level with the most susceptible line Heinz 1706. NC 1CELBR represented a line with a resistance level between these two lines (Table 1, Fig. 1).

**Fig 1.**
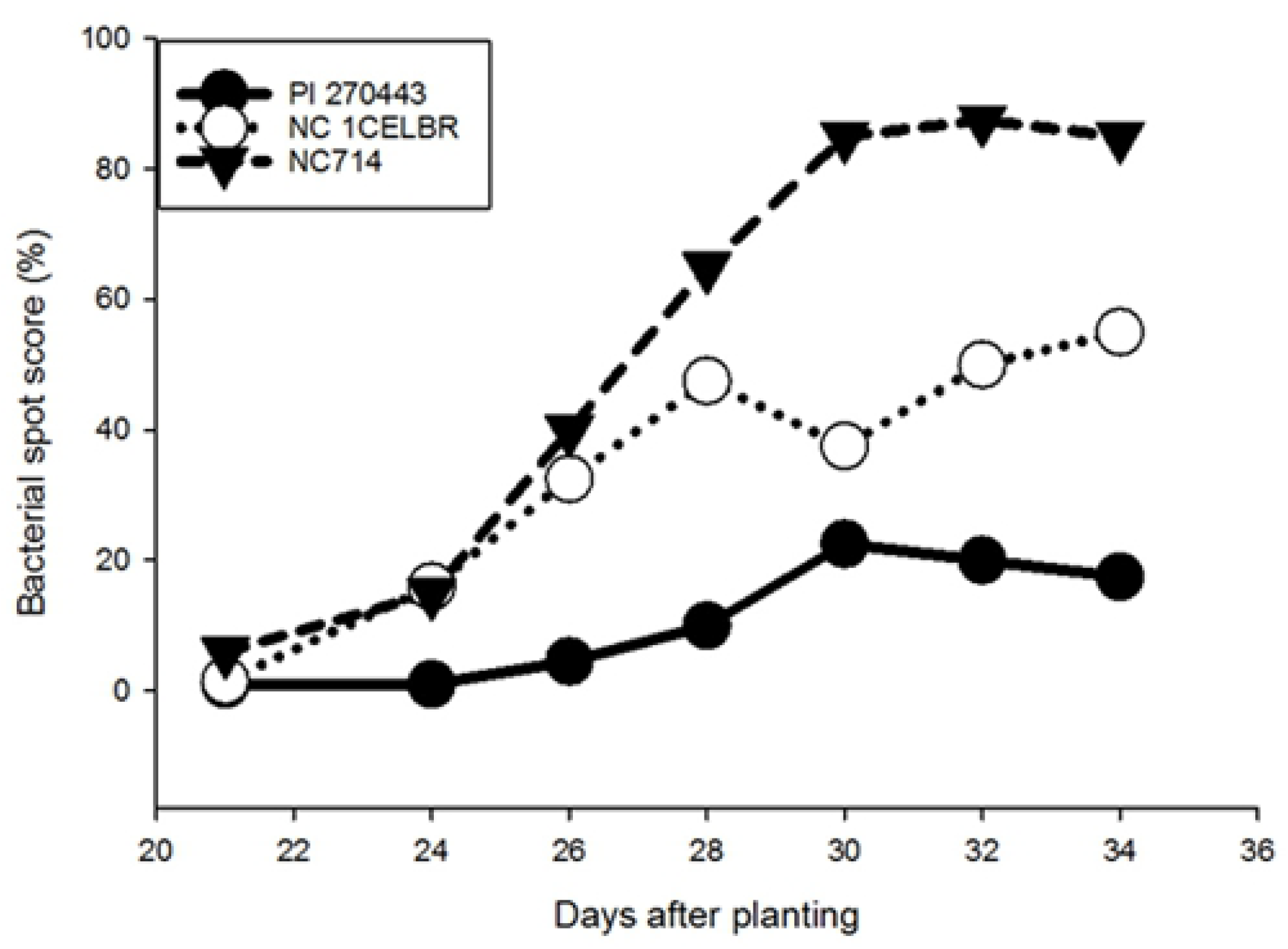
Resistance level of three selected tomato lines for BS.

### RNA-Seq data analysis showing gene expression variation between tomato breeding lines was much more significant than treatment

To investigate the differences in transcriptome associated with inoculation of BS race T4, 12 RNA-Seq libraries were constructed and sequenced with two biological replicates for three tomato lines inoculated with BS (IN) or without BS inoculation (CK). A total of 267 million reads were generated for these RNA-Seq libraries. These reads were processed to remove low quality reads and then mapped to Heinz 1706 genome assembly SL3.0. The mapped RNA-Seq reads were quality filtered and resulted in a total of 234 million mapped reads (at least 6 million per library) for subsequent bioinformatics analysis. The data of these RNA-Seq libraries have been deposited in NCBI’s Gene Expression Omnibus [41] and are accessible through GEO Series accession number GSE135232.

The first step of our analysis was to check the relationship of all RNA-Seq libraries sequence using plotMDS function of Edge R package [34]. Ideally, replicated samples from the same group should cluster together in the plot, while samples from different sample groups form separate clusters [34]. In this plot, we found that samples of each tomato lines are grouped, although RNA-Seq libraries for uninoculated (CK) and inoculated (IN) samples separated slightly from each other for the same tomato line (Fig. 2). This plot pattern indicated that gene expression variation between tomato breeding lines was more significant than that induced by BS inoculation treatment, i.e., the difference between different tomato lines was much bigger than that between control and inoculation within the same tomato line. This might result from the inoculation method. We inoculated the breeding lines by spray method, which was less efficient compared to vacuum infiltration method. Therefore, fewer tomato cells were exposed to the pathogen and thus their gene expression was less represented. Because of these possible variations level, we adopted the strategy of analyzing genes with expression changes between control and inoculated samples within the same genotype, and then compare inoculation induced DEGs between different tomato lines.

**Fig 2.**
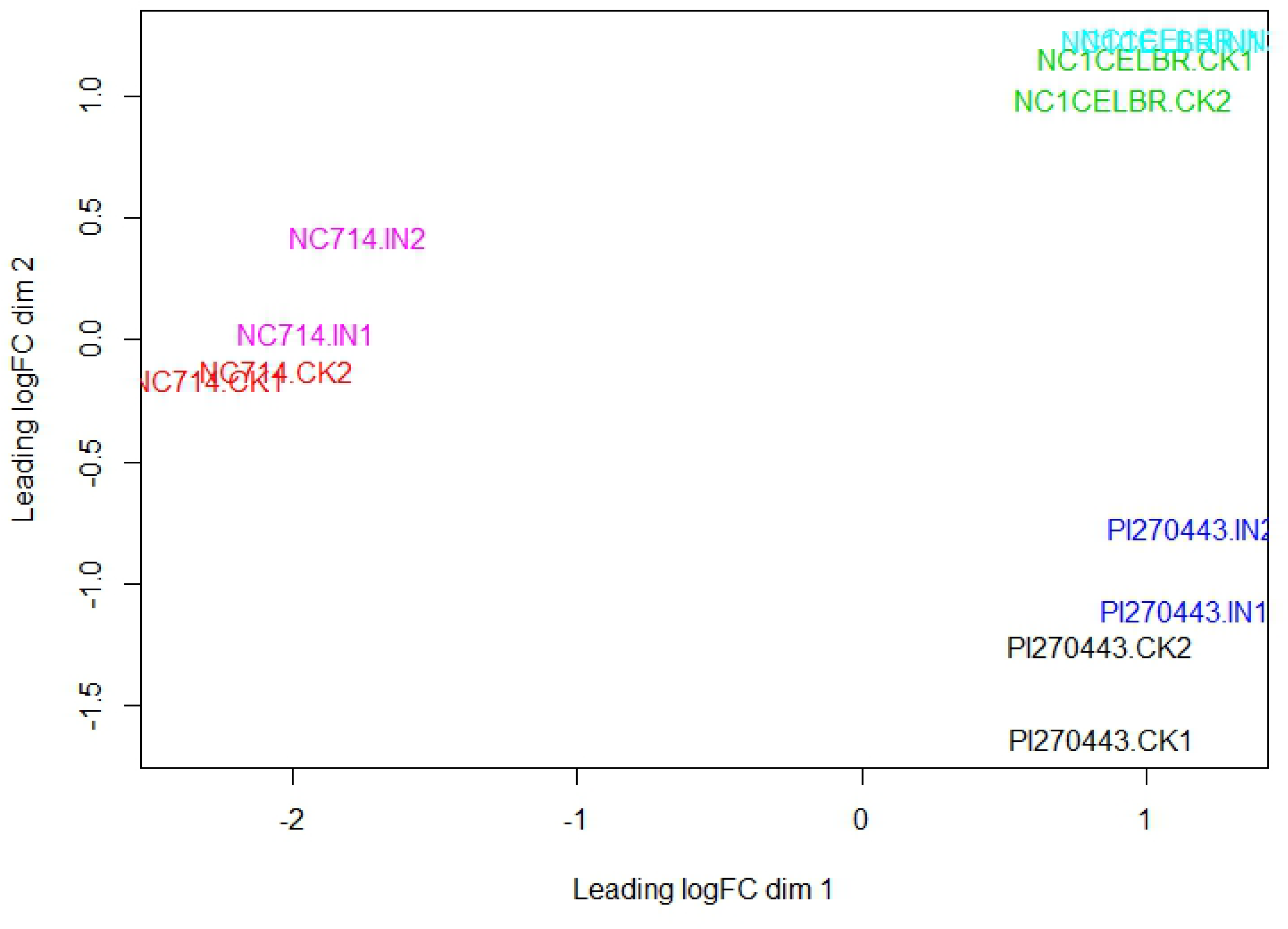
Multidimensional scaling plot for the relationship of each RNA-Seq library. Distance between libraries was calculated as the leading log fold change for biological variation between libraries [42]. The label of “IN” represent BS inoculated samples, and “CK” represents a control sample without inoculation of BS.

### DEG analysis showing resistant tomato line PI 270443 reacts to inoculation of BS is stronger than other lines

Differentially expressed genes (DEGs) were screened between control and inoculated tomatoes for each breeding lines based on 2-times fold change of transcript abundance and FDR<0.05. A total of 1,362 differentially expressed genes were identified from this study. Among them, PI 270443 had 834 (346 up-regulated, and 488 down-regulated) genes, NC 1CELBR had 154 (71 up-regulated, 83 down-regulated), and NC 714 had 373 (93 up-regulated, 277 down-regulated) DEGs between control and inoculated samples, respectively (Fig. 3A, Table S1).

**Fig 3.**
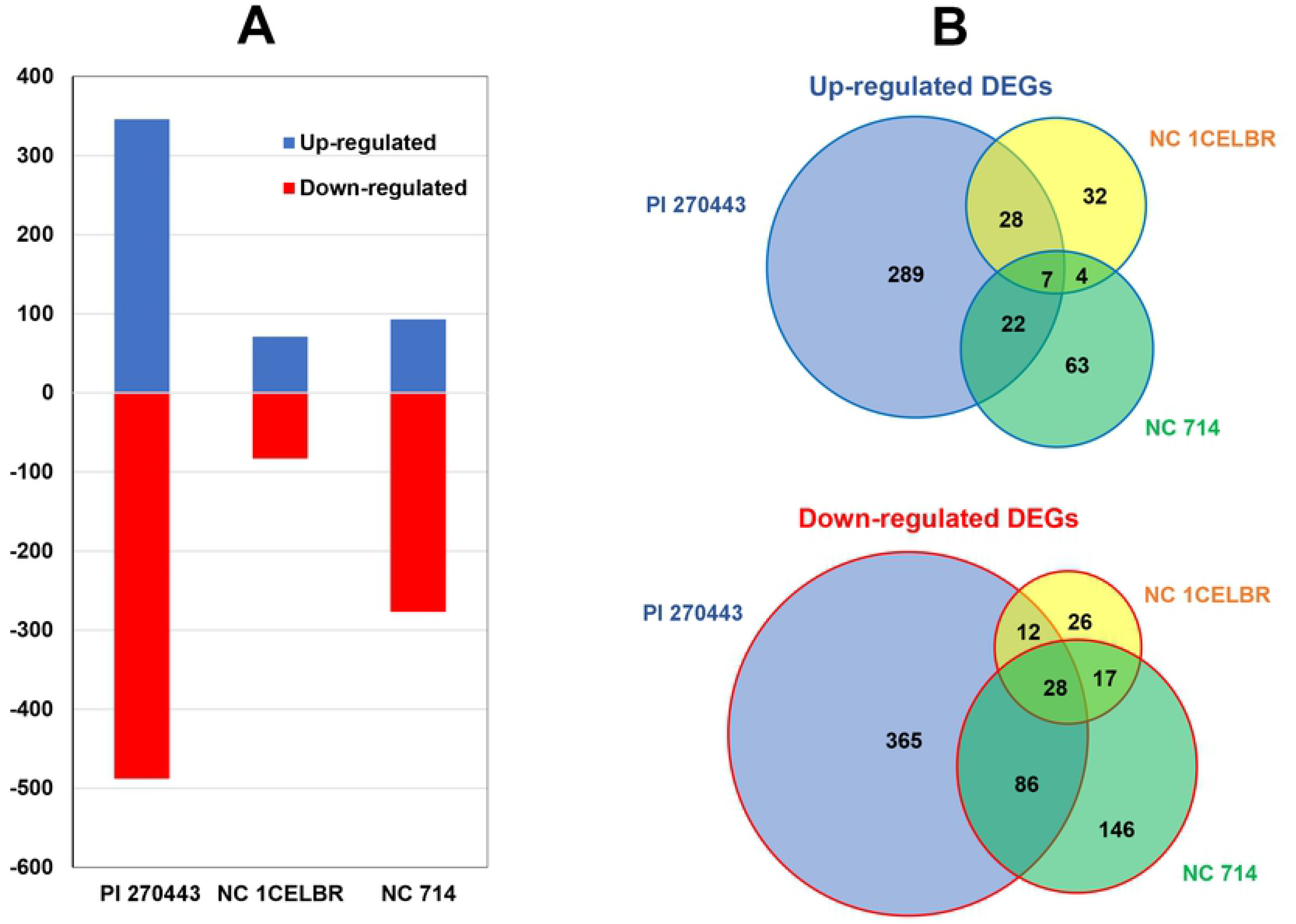
DEGs in three tomato lines induced by BS inoculation. (A) The number of up-regulated (blue bar) and down-regulated DEGs (red bar, with the label of “-”) in tomato genotypes with respect to the inoculation of *X. perforans* race T4. (B) Venn diagram showing the up-regulated (blue outlined) and down-regulated (red outlined) differentially expressed genes (DEGs) in three tomato lines PI 270443, NC 1CELBR, and NC 714. An entire list of differentially expressed genes (DEGs) in all three tomato breeding lines is presented as supplementary material (Table S1).

Overall, there were 510 up-regulated genes and 852 down-regulated genes. Among them, only 35 genes were common (seven up-regulated and 28 down-regulated) among all three tomato lines, associated with bacterial spot (BS) resistance. There were only 112 DEGs between PI 270443 and NC 714, whereas there were only 42 up-regulated DEGs between PI 270443 and NC 1CELBR. There were 21 genes up-regulated DEGs between NC 1CELBR and NC 714 (Fig. 3B).

### More GO terms were enriched for up-regulated DEGs in resistant PI 270443, whereas more GO terms were enriched for down-regulated DEGs in susceptible NC 714

Functional analysis of DEGs for different tomato lines conducted based on gene ontology (GO) enrichment analysis using AgriGO based on tomato ITAG3.2 version annotation and background. We expected that some GO terms were enriched (over-represented) after BS inoculation, and provide clues to understand the reaction of BS inoculation for different tomato lines. Based on this analysis, we found more GO terms enriched for up-regulated DEGs in resistant PI 270443 line, and more GO terms were enriched for down-regulated DEGs in susceptible NC 714 line (Fig 4, Table S2).

**Fig 4.**
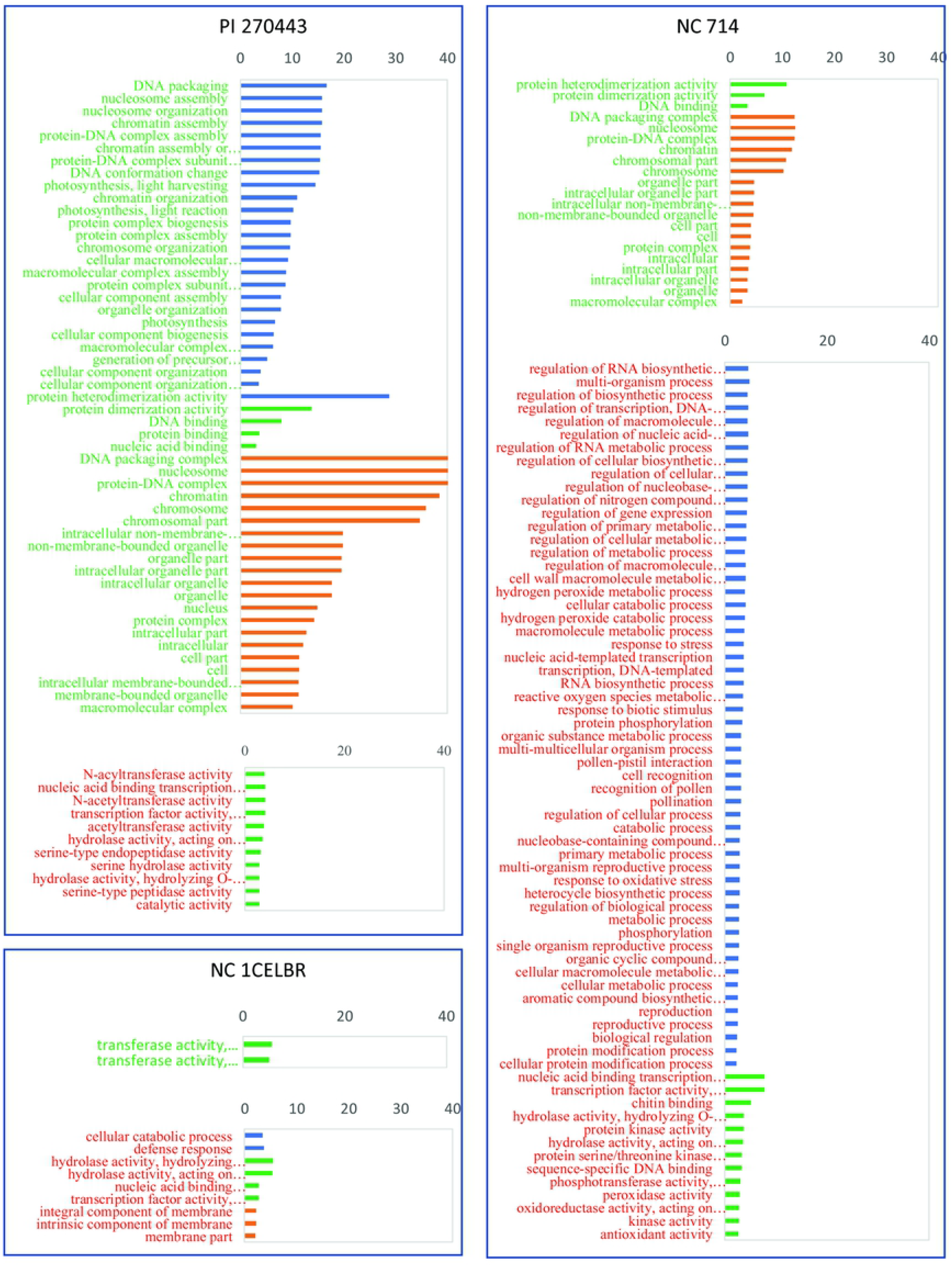
GO term enriched for DEGs in different tomato lines after BS inoculation. Labels in green and red represent up-regulated and down-regulated DEGs, respectively. Bars colored in blue, green and orange represent biological process, cellular component and molecular function GO category, respectively. Scale is log10 (1/p-value).

In detail, there were 51 GO terms enriched for 346 up-regulated DEGs from PI 270443. Among the enriched GO terms, the important GO terms included protein binding, cell parts, cell, intracellular, intracellular part, intracellular organelle, organelle, nucleic acid binding, DNA binding, and protein complex (Table S2). Additional important GO enriched terms were intracellular non-membrane-bounded organelle, non-membrane-bounded organelle, and macromolecular complex among others.

For 488 down-regulated DEGs from PI 270443, there were 11 GO enriched terms with a relatively low level of enrichment, and all of them had molecular functions. The important GO terms were catalytic activity, nucleic acid binding transcription factor activity, transcription factor activity, sequence-specific DNA binding, hydrolase activity, acting on glycosyl bonds, hydrolase activity, hydrolyzing O-glycosyl compounds, serine hydrolase activity, and serine-type peptidase activity among others (Fig 4, Table S2).

There were only two GO terms enriched for NC 1CELBR for 71 up-regulated DEGs, at the level of 4, which are transferase activity, transferring hexosyl groups and transferase activity, transferring glycosyl groups. Nine GO terms were enriched for 81 down-regulated DEGs from NC 1CELBR, including membrane part, hydrolase activity, hydrolyzing O-glycosyl compounds, hydrolase activity (acting on glycosyl bonds, an integral component of the membrane), cellular catabolic process, and defense response (Fig 4, Table S2).

For 96 up-regulated DEGs from NC 714, there were 21 GO enriched terms. The significantly enriched terms are associated with cellular components such as DNA and chromosome (Table 6). For 277 down-regulated DEG for NC 714, there were 67 GO terms enriched, which include response to stress (GO:0006950) and biotic stimulus (GO:0009607) among others, including protein kinase activity, peroxidase activity, antioxidant activity, response to biotic stimulus, cell recognition, chitin-binding, protein phosphorylation, regulation of cellular process, cellular metabolic process, and response to oxidative stress among others (Fig 4, Table S2).

Overall, there were several enrichments of GO terms for up-regulated DEGs from PI 270443, which might be associated with the resistance of this genotype. Contrary to this, more enriched GO terms for down-regulated DEG of NC 714 might be a clue to the susceptibility of NC 714. It should be noted that there was no overlap GO terms for all up-regulated DEGs from these three lines, but all 21 GO terms enriched in up-regulated DEGs of NC 714 were overlapped with GO enrichments from PI 270443. As to down-regulated DEGs, there were four GO terms enriched for down-regulated DEGs for all these three lines, which might refer to a common reaction to BS. These GO terms include nucleic acid binding transcription factor activity, transcription factor activity, sequence-specific DNA binding hydrolase activity, and acting on glycosyl bonds hydrolase activity, hydrolyzing O-glycosyl compounds.

### More up-regulated DEGs were identified for resistant PI 270443 and more down-regulated DEGs found for susceptible NC 714 in the biotic stress pathway

To understand the responses of these tomato lines to BS infection, MapMan was adopted to classify DEGs of these tomato lines based on the more comprehensive annotation of tomato genes as described in the methods section. This analysis was conducted to decipher the involvement of DEGs in various cellular processes using the MapMan software. In Fig 5, we present the distribution of DEGs in biotic stress pathway, which is more associated with disease resistance and development.

**Fig 5.**
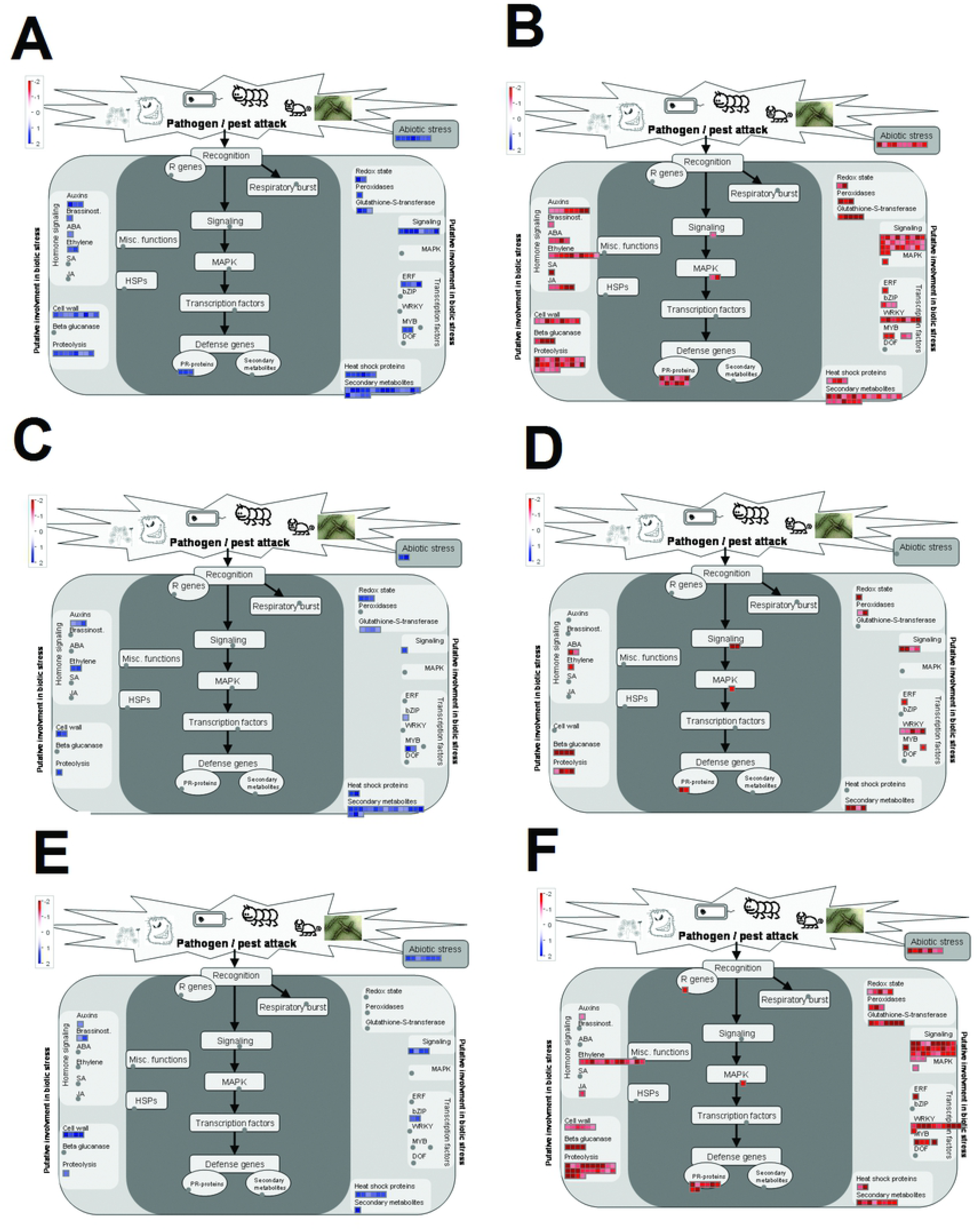
BS inoculation induced DEGs in the biotic pathway for different tomato lines. DEGs identified in biotic stress pathway for PI 270443, NC 1CELBR, and NC 714. Left panels represent up-regulated DEGs (blue color), and right panels are down-regulated DEGs (red color).

Overall, highly resistant line PI 270443 had more up-regulated DEGs (67) in the biotic stress pathway, followed by 34 up-regulated DEG in NC 1CELBR, and 21 up-regulated DEGs in NC 714. On the other hand, up-regulated DEGs in PI 270443 had more bins (catalogs defined for genes relating functional processes in MapMan) and with more DEGs in each bin as well. For instance, PI 270443 had 3 up-regulated PR-protein genes in the core of biotic stress pathway (dark area in Fig 5) and had DEGs associated with ABA and peroxidase, which were not present in other two lines. There were also more DEGs associated cell wall, proteolysis, signaling, ERF, and secondary metabolites for PI 270443 than that in the other two lines. Although overall DEGs and enriched GO term for NC 1CELBR seemed under-estimated in our analysis, this medium resistant tomato line still had more up-regulated DEGs in biotic stress pathway than that of NC 714. The DEGs were associated with Auxins, Ethylene, Redox associated genes, Glutathione-S-transferase, heat shock proteins, and secondary metabolites.

As to down-regulated DEGs in this biotic stress pathway, PI 270443 had 154 DEGs, followed by 126 in NC 714, and 33 in NC 1CELBR. As to each catalog, the pattern of PI 270443 and NC 714 was similar in number, but NC 1CELBR had few DEGs in each bin. One noticeable pattern was that one DEG (solyc12g097000) defined as R gene which is important for disease resistance was down-regulated in NC 714, and no such DEG was found in biotic stress pathway for other two lines.

**Table 2:** Summary of RNA-Seq library information.

| Sample ID | No. of raw reads | No. of clean reads | No. of reads mapped to the genome | No. of reads mapped to genome after filtering |
| --- | --- | --- | --- | --- |
| PI270443-CK1 | 17,401,320 | 17,366,776 | 15,679,410 | 14,493,893 |
| PI270443-CK2 | 16,652,433 | 16,627,277 | 14,877,660 | 14,493,893 |
| NCCELBR1-CK1 | 22,569,408 | 22,539,261 | 20,742,997 | 20,134,965 |
| NCCELBR1-CK2 | 15,898,554 | 15,872,147 | 13,799,117 | 13,418,896 |
| NC 714 -CK1 | 16,716,909 | 16,672,953 | 15,014,160 | 14,652,561 |
| NC 714 -CK2 | 8,045,470 | 8,028,488 | 6,783,492 | 6,615,839 |
| PI270443-IN1 | 15,187,278 | 15,150,128 | 12,830,082 | 12,458,547 |
| PI270443-IN2 | 10,417,144 | 10,390,478 | 8,475,406 | 8,204,475 |
| NCCELBR1-IN1 | 26,100,659 | 26,015,053 | 24,082,176 | 23,340,952 |
| NCCELBR1-IN2 | 37,792,194 | 37,721,275 | 35,177,779 | 34,073,612 |
| NC 714 -IN1 | 57,979,118 | 57,873,646 | 54,422,298 | 52,813,125 |
| NC 714 -IN2 | 22,812,966 | 22,762,930 | 19,933,275 | 19,364,162 |

Table 3 presents the overlapping DEGs in biotic stress pathway for these tomato lines. It shows that among 67 DEGs up-regulated in PI 270443, only two were common up-regulated DEGs in all three lines, which might be associated with common reaction to the inoculation of BS race T4. There were 16 DEGs up-regulated in both PI 270443 and NC 1CELBR, which are resistant to BS. One noticeable feature was that half of them are involved with secondary metabolism of flavonoids. Furthermore, *CCoAOMT* gene, which was reported to be disease-resistance associated in maize [43–49], was found as the common up-regulated DEG in these two resistant tomato lines. As to up-regulated DEGs specific for PI 270443 only, some group of genes was overrepresented, including those associated with cell wall cellulose synthesis, APETALA2/Ethylene-responsive element binding protein family, protein degradation related, and stress-related genes, etc.

**Table 3.**
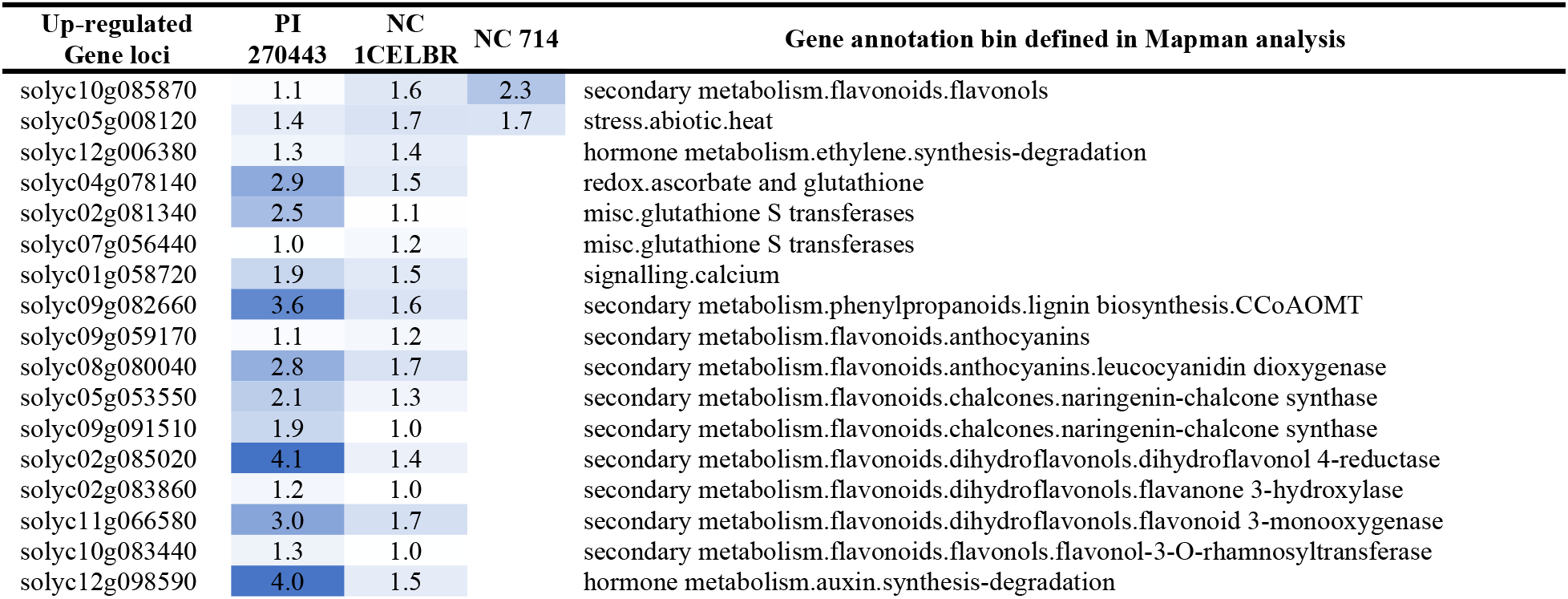
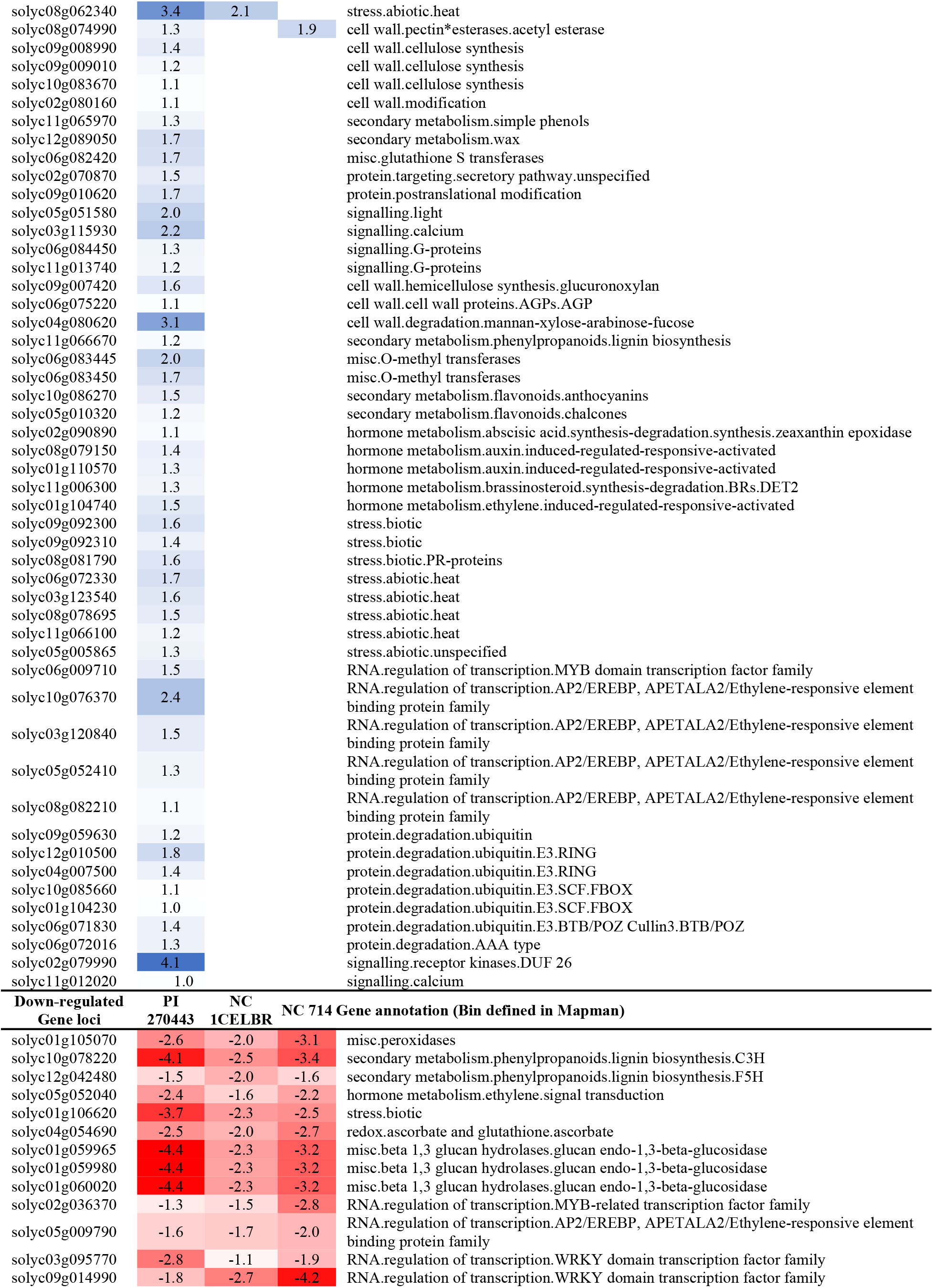
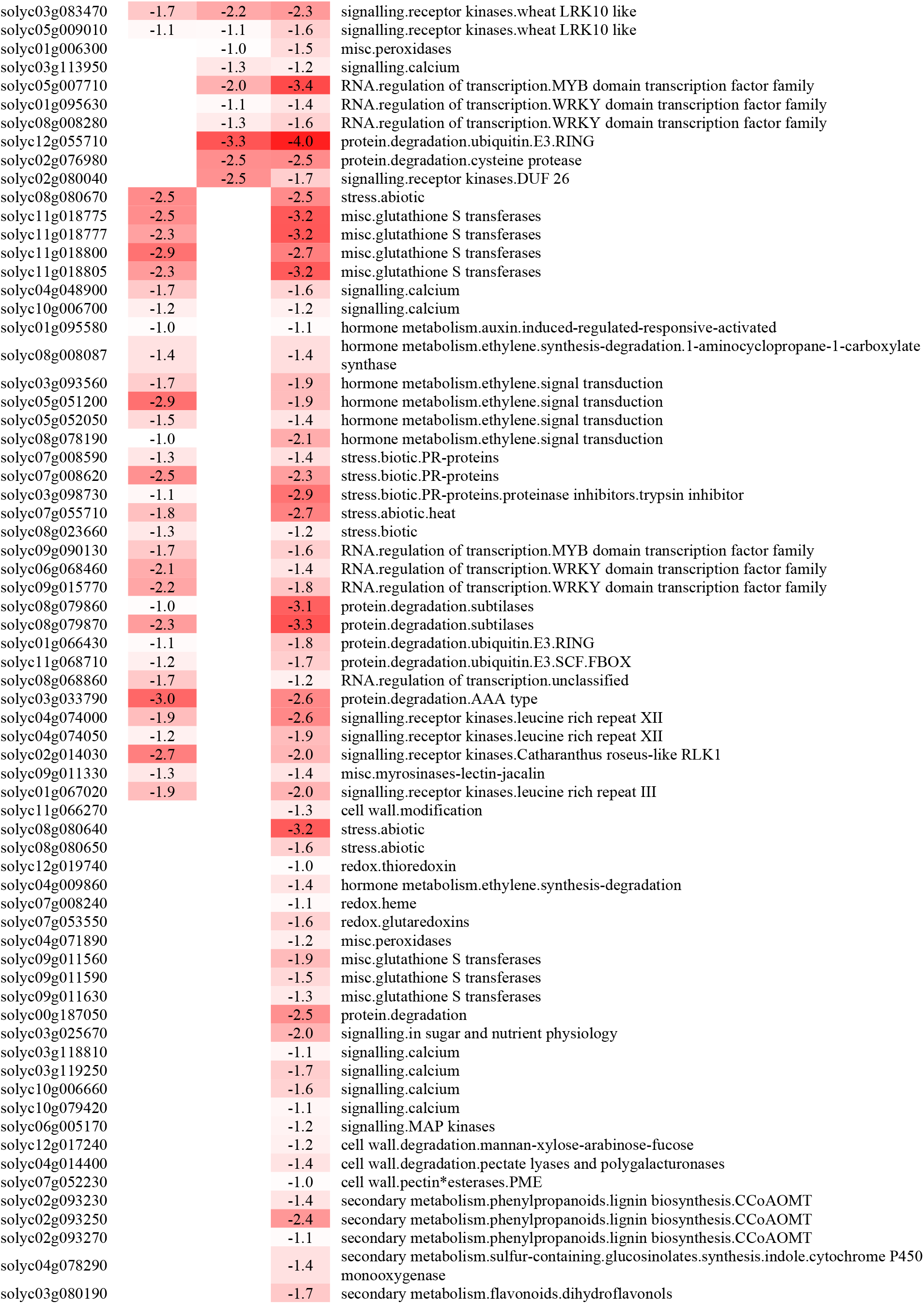
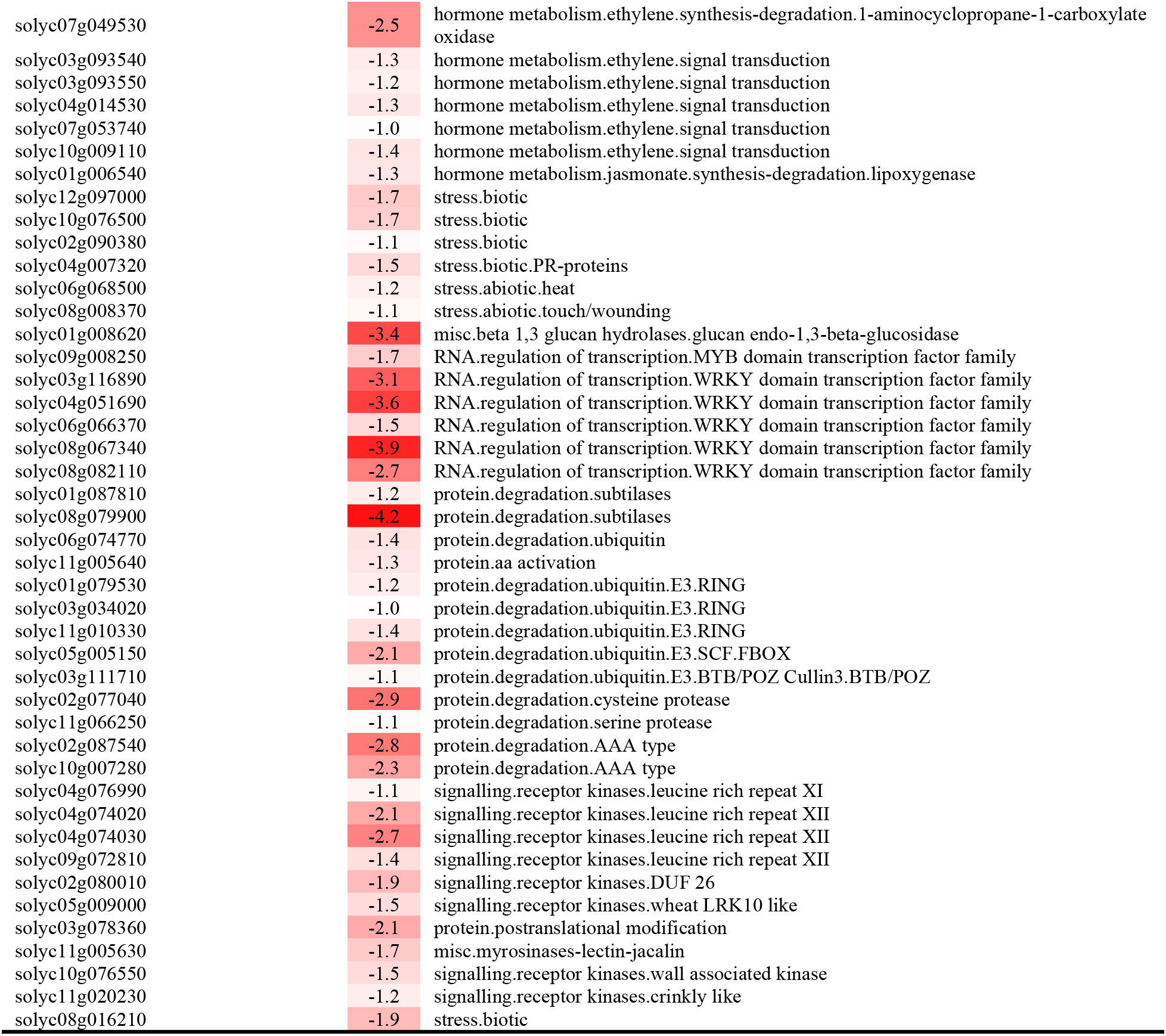
DEGs in biotic stress pathway associated with resistance and susceptibility. The upper part of the table list all up-regulated biotic stress pathway DEGs in the resistant line PI 270443 with expression change (log2) between inoculated and control samples, and lower part of the table list all down-regulated biotic stress pathway DEGs in the susceptible line NC 714.

Among 125 down-regulated DEGs in NC 714, which might be associated with susceptibility to BS race T4, there were 15 commonly down-regulated genes in all three lines, including two secondary metabolisms lignin genes *C3H* and *F5H*, and three genes encoding *β*-1,3 glucan endo-1,3-*β*-glucosidase. These DEGs, we believe, are common reactions to the inoculation of BS. Down-regulated DEGs specific to NC 714 included three *CCoAOMT* genes, five WRKY domain transcription factor genes, and five genes related to protein degradation/ubiquitin in the list of the common down-regulated gene for PI 270443 and NC 714 (Table 3).

### Sequence variation called from transcriptome identify the relationship of chromosome with resistance

SNP molecular marker analysis was conducted based on a sequence of RNA-Seq reads mapped to reference genome sequence of tomato Heinz 1706 (SL3.0), which is the most susceptible to BS race T4. Overall a total of 23,253 SNP/INDELs site were called in PI 270443, NC 1CELBR, and NC 714 against Heinz 1706. Among these SNP/INDELs, 18,238 could be located within the sequence of 6,676 gene models (annotation version ITAG3.2). Distribution of these SNP/INDELs and associated genes on the individual chromosome are listed in Table 4. As SNP/INDELs or associated genes distribution in individual tomato line, these were defined as a homo form of alternative sequence to Heinz 1706, i.e., GT:1/1 in genotype section of VAR file. More detailed SNP/INDELs raw information can also be found in Table S3.

**Table 4:** Distribution of SNP/INDELs on different chromosomes of tomato.

| Chr | SNP/INDELs loci (Gene with SNP/INDELs in parenthesis) |  |  |  |
| --- | --- | --- | --- | --- |
|  | Total | PI 270443 | NC 1CELBR | NC 714 |
| 1 | 1850 (728) | 781 (274) | 734 (248) | 1219 (481) |
| 2 | 1285 (531) | 437 (167) | 563 (191) | 875 (397) |
| 3 | 2162 (720) | 595 (230) | 739 (247) | 1575 (584) |
| 4 | 3306 (825) | 1143 (298) | 2061 (488) | 1826 (556) |
| 5 | 1266 (455) | 633 (168) | 544 (161) | 958 (376) |
| 6 | 1149 (432) | 414 (125) | 378 (110) | 949 (379) |
| 7 | 1242 (366) | 447 (146) | 398 (143) | 937(309) |
| 8 | 1359 (466) | 517 (139) | 559 (152) | 1036 (400) |
| 9 | 2210 (535) | 675 (200) | 1522 (345) | 1264 (397) |
| 10 | 1869 (473) | 825 (242) | 1244 (289) | 904 (273) |
| 11 | **3835 (716)** | **2893 (544)** | **2894 (545)** | **1699 (341)** |
| 12 | **1223 (377)** | **944 (311)** | **938 (301)** | **488 (152)** |
| Total | 23253 (6676) | 10666 (2865) | 12942 (3239) | 14063 (4667) |

These results showed an interesting phenomenon that in NC 714, the most susceptible tomato breeding line had more SNP/INDELs against reference genome of similar susceptible Heinz 1706’s, whereas the most resistant PI 270443, which is expected to be different from Heinz 1706 (susceptible to BS), had fewer number of SNP/INDELs. The situation of the individual chromosome was different. For instance, chr11 and chr 12 of NC 714 had relatively less SNP/INDELs and associated genes against Heinz 1706, but PI 270443 and NC 1CELBR had more SNP/INDELs and associated genes, respectively.

On further evaluation of distribution of sequence variation as shown in the locations of genes with SNP/INDELs on each chromosome (Fig 6A), we noticed that the patterns of genes with SNP/INDELs on chr11 SNP/INDELs from PI 270443 and NC 1CELBR were almost the same, which can be further confirmed by comparison of SNP/INDELs between these two lines resulting in over 95% identical on chr11 (Fig 6B). One key insight of this pattern was that chr11 has resistance QTLs as identified in the previous study, and these QTLs contribute 25% resistance to race T4 [11]. This pattern suggested that QTLs on chr11 might contribute to resistance of PI 270443 and NC 1CELBR.

**Fig 6.**
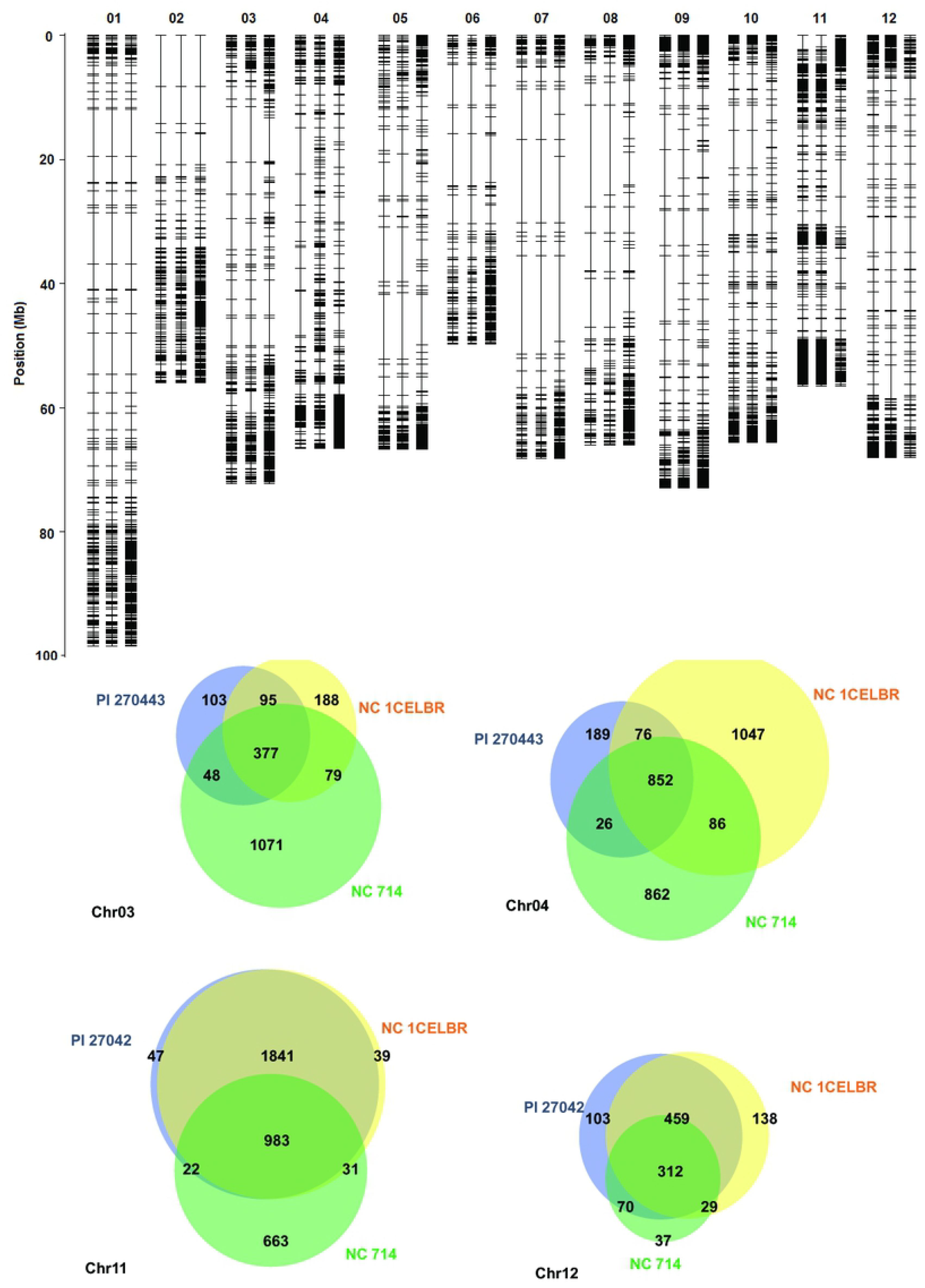
Analysis of sequence variation identified from transcriptomes of three tomato lines. (A) Distribution of genes with SNP/INDELs on different chromosomes. (B) Venn diagram for overlaying SNP/INDELs identified from individual tomato lines for chromosome 3, 4, 11, and 12 (labeled as chr03, chr04, chr11, and chr12).

In addition to similar sequence variation pattern on chr11, which has resistance QTL as reported before [11], these two lines also share some common sequence variation on chr 12 (Fig 6B), which has QTLs related to susceptibility to race T4 of BS [11]. Therefore, such common sequence variation in PI 270443 and NC 1CELBR against the reference genome of susceptible Heinz 1706 might be associated with reduced susceptible QTLs to the BS. SNP/INDELs on other chromosomes in these three lines were more diverse, such as chr 3 and chr 4 as shown in Fig 6B, even though it was reported that chr 3 has resistance QTL to BS [11].

## DISCUSSION

### Bacterial spot inoculation experiments and genotype represented for RNA-Seq analysis

The primary objective of this study was to determine and identify the unique DEG in response to the BS so that we can identify the specific genes associated with BS resistance in tomato. It has been a challenge to determine the resistance genes hence to introgress the resistance into the breeding lines with desirable fruit quality in tomato despite a lot of efforts towards the identification of QTL associated with BS resistance. In this study, the response of tomato genotypes to the BS race T4 was found as expected. For example, PI 114490, which is resistant to all races of *Xanthomonas*, also had less disease severity.

On the other hand, NC 714, which does not have any disease resistance to any race of *Xanthomonas*, was found to develop severe disease. An exciting aspect of this study was that PI 270443 had less disease than even in PI 114490. In several papers, PI 114490 has been reported to have resistance to all available races [10, 11, 50-53]. In our study, however, a genotype more resistant than PI 114490 is available and whatever genes identified from the line may provide more robust information towards the identification of resistance genes. It should be noted that the race of BS reported from NC is race T4 [22]. This indicated that the source of resistance might be associated with a single race. This will still be a beneficial material to address this critical issue. Race T4 is widely distributed in FL and NC tomato growing regions [22].

### RNA-Seq analysis

RNA-Seq analysis successfully identified DEGs associated with critical biological and cellular processes. While some of the genes reaction was unique to the *Xanthomonas* race T4, others were already reported either with plant defense system or with stress tolerance. These are the types of genes related to cellular and biological functions. In most of the gene expression or RNA-Seq studies, many pathways are involved, although biotic stress pathways were especially focused on specific disease resistance-associated genes.

As expected, there was a significant difference between inoculated and control breeding lines for DEGs when evaluated based on RNA-Seq analysis. Du, Wang (13] performed the gene expression analysis using race T3 of *X. perforans* in tomato and identified more DEG in resistant line PI 114490 than in susceptible line OH88119, and identified different sets of genes associated with cellular and molecular processes. Our analysis revealed that resistant line PI 270443 did involve with more DEGs overall compared to susceptible line NC 714, but NC 1CELBR with a medium level of resistance has fewer DEGs induced by inoculation of BS race T4. It might be due to inoculation method as described above. However, when we closely evaluated the up-regulated DEGs in biotic stress pathway, we can identify more biotic stress associated DEGs in NC 1CELBR than in NC 714, even though total up-regulated DEGs or GO term enriched in NC 1CELBR were much less than that in NC 714.

All identified DEGs would be valuable resources for the evaluation of plant reaction to BS infection, and GO term enrichment is the most adopted strategy to show overall response based in term of over-representation. However, in this study, we also adopted MapMan based analysis to identify genes associated with disease resistance. Our result showed a noticeable feature that there were a higher number of up-regulated genes induced by the inoculation of BS with the resistant tomato line, and a larger number of down-regulated genes found in the susceptible line after *X. perforans* race T4 inoculation. We tend to think that such patterns would provide us a good chance to find resistance-related genes in resistant genotype and genes associated with disease development in susceptible genotype. For instance, in the core genes of biotic stress pathway, we found that only resistant line PI 270443 had three up-regulated PR genes (loci solyc09g092300, solyc08g081790, and solyc09g092310) associated with inoculation of BS race T4. On other and, only susceptible NC 714 line has a down-regulated R gene (locus solyc12g097000) among three lines. These genes were not reported in the study conducted by Du, Wang (13]. A possible reason is that we analyzed samples collected two days after inoculation (48 hai), which is different from the condition of samples collected by Du, Wang (13] after 6 hai and 6 dai. Additionally, the genetic background of their especially susceptible variety is much different from the tomato lines we analyzed.

As to other bins of putative genes involved in biotic stress pathway, we identified genes for Glycosyl hydrolases. Glycosyl hydrolases (GHs) comprise a large assembly of enzymes that hydrolyze the glycosidic bond between carbohydrates or between carbohydrates and non-carbohydrate moieties. GHs are grouped into various families based on amino acid sequence similarities. These proteins perform diverse functions in both plants and microbes. Many pathogenesis-related (PR) proteins belong to the GH group. GH family 17 and families 18 and 19, which contain β-1,3 glucanases and chitinases, respectively, form an important part of the defense arsenal of plants against fungal pathogens [54–56].

We also found many genes associated with secondary metabolites involved in the reaction of inoculation of BS. For instance, in addition to many up-regulated genes for secondary metabolites of flavonoids, we identified *CCoAMOT* (solyc09g082660) was up-regulated in PI 270443 after inoculation, and several *CCoAMOTs* (solyc02g093230, solyc02g093250, and solyc02g093270) were down-regulated in NC 714 only. It should be mentioned that CCoAMOT has been reported as a gene involved in disease resistance [56–58]. On the other hand, *C3H* (solyc10g078220) and *F5H* (solyc12g042480) for lignin biosynthesis were down-regulated in all tomato genotypes, which might be a common reaction to the inoculation of BS.

Genes with regulation functions are more complex to analyze due to too many associated genes involved, but we still found that after inoculation of BS race T4, there are more ethylene-responsive transcript factor genes up-regulated in resistant PI 270443, and many *WRKY*s, on another hand, were down-regulated in susceptible NC 714. Such reaction seems to be different from that reported by Du, Wang (13], since *WRKY*s including Solyc03g116890, Solyc04g051690, Solyc06g066370 and Solyc08g082110 were up-regulated in both resistance line PI 114490 and susceptible line OH 88119 after 6 hai, but down-regulated 6 dai in these two lines in the BS race T3 inoculation experiment. It is possibly due to the different inoculation time of tomato samples with BS.

### SNP/INDELs analysis

Although DEGs analysis can reveal overall functions specific to biotic stress pathway for genes associated with disease resistance and susceptibility processes, sequence variation within expressed genes can also contribute to disease resistance even if they are not DEGs. Sequence variation in gene coding region can alter the function of genes, and thus may affect disease resistance. For instance, the sequence variation in *Rx4* was found to be associated with the resistance to race T3 of BS in tomato [3]. Therefore, we have taken advantage of transcriptome information in this study to call SNP/INDELs and hope to find genes with sequence variation between tomato lines resistant and susceptible to BS.

Our results showed more SNP/INDELs in NC 714 against Heinz 1706, suggesting a more diverse genetic background between these two BS susceptible lines. However, such sequence variation on chr11 was small, which suggested that both lines lack resistance genes to BS. The chr11 of PI 270443 and NC 1CELBR were identical based on SNP/INDELs, which was different from that of NC 714 and Heinz 1706. This phenomenon suggested that the chr11 contributes to the resistance of tomato lines to BS in a similar way. For instance, chr11 was reported to have QTLs contributing ~(*R^2^*=29.4%) resistance to BS [11]. Also, chr 3 ~(*R^2^*=4.3%), which has been reported with resistance elements to BS, did not show such pattern for these two resistant tomato lines and might explain the different resistance level for PI 270443 and NC 1CEBLR to BS infection. To be more focused on resistance gene to BS race T4, we feel that it would be easy to specially concentrate on chr11 to find the genes associated with resistant QTLs, and listed all DEGs and genes with sequence variation in biotic stress pathway to identify genes similar to reported resistance genes.

Line PI 128216 derived from *S. pimpinellifolium*, has been reported to carry the gene *Rx4* located on chromosome 11 conferring hypersensitivity and field resistance to race T3. An NBS-LRR class of resistance gene was fine-mapped near 45.1 kb region between pcc17 and pcc14 molecular markers. Six SNP and one INDEL molecular markers were also developed from this region, which was found to be useful for the MAS of this gene [3].

The recessive genes, *rx1* and *rx2*, conferring resistance to race T1, were derived from Hawaii 7998 and mapped onto chr 1, whereas *Rx3* was mapped onto chr3 [4]. Modifying susceptible alleles have been reported from chr 3, 5, 9 and 11. The genes *avrXv3, Xv3/ avrXv3 and Xv3/ Rx4* derived from Hawaii 7981 and PI 126932, respectively were located on chr11, conferring resistance to race T3 [59, 60].

By blasting published cloned *Rx4* candidate transcript sequence for *S. lycopersicum* cultivar Hawaii 7981, OH88119 and *S. pimpinelifolium* PI 128216 (JF743044.1, JF743043.1, and JF743045.1, respectively), we found that the candidate *Rx4* gene is a homolog of solyc11g069020, which has no SNP/INDELs identified in this study due to their low expression levels in the tissues tested. On the other hand, we can find multiple PR genes around this *Rx4* candidate gene (<2Mb, Table 5) and these PR genes have SNP/INDLEs specific in the resistant tomato lines PI 270443 and medium resistant line NC 1CELBR against susceptible line NC 714 and Heinz 1706 (Table 5). Although these PR genes are not identified as BS induced DEGs for these two lines, they might still function diversely from their homologs in susceptible tomato and might be candidate genes function as multiple loci for resistance to BS race T4.

**Table 5.**
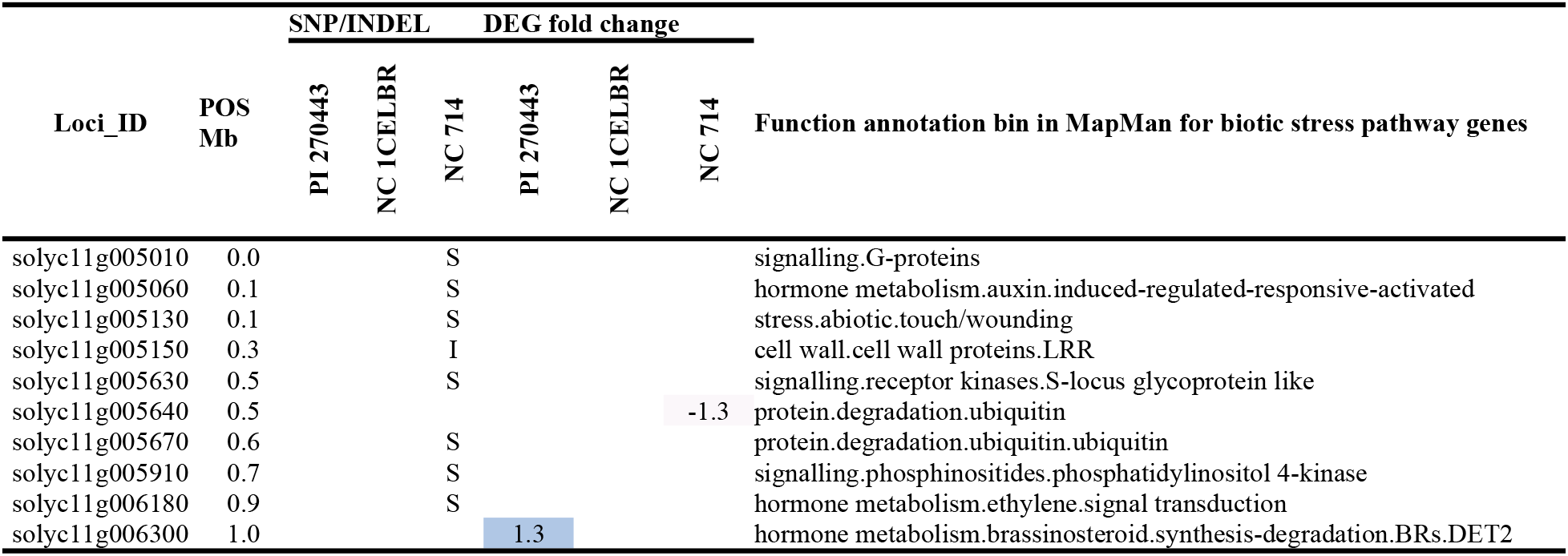
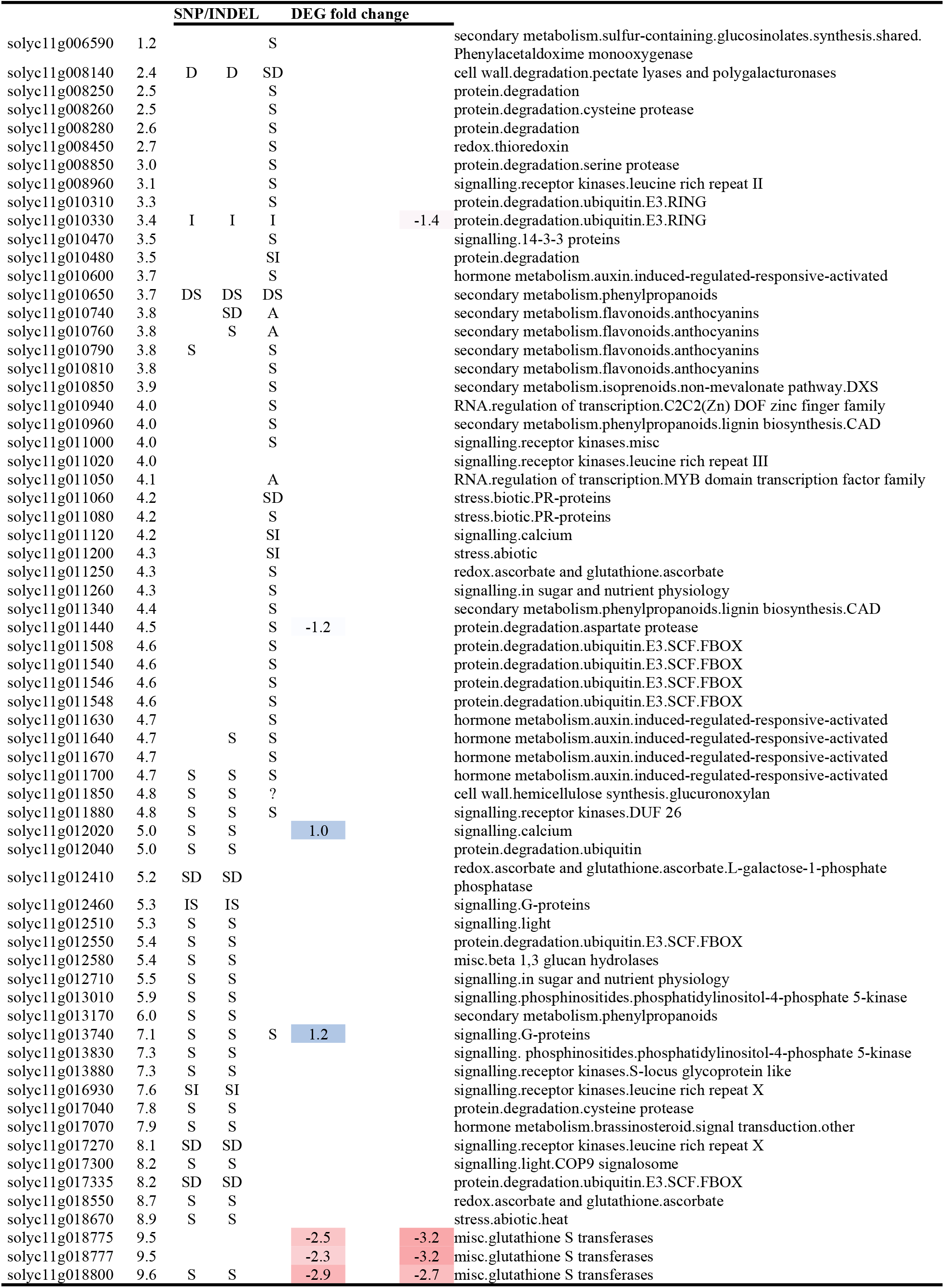
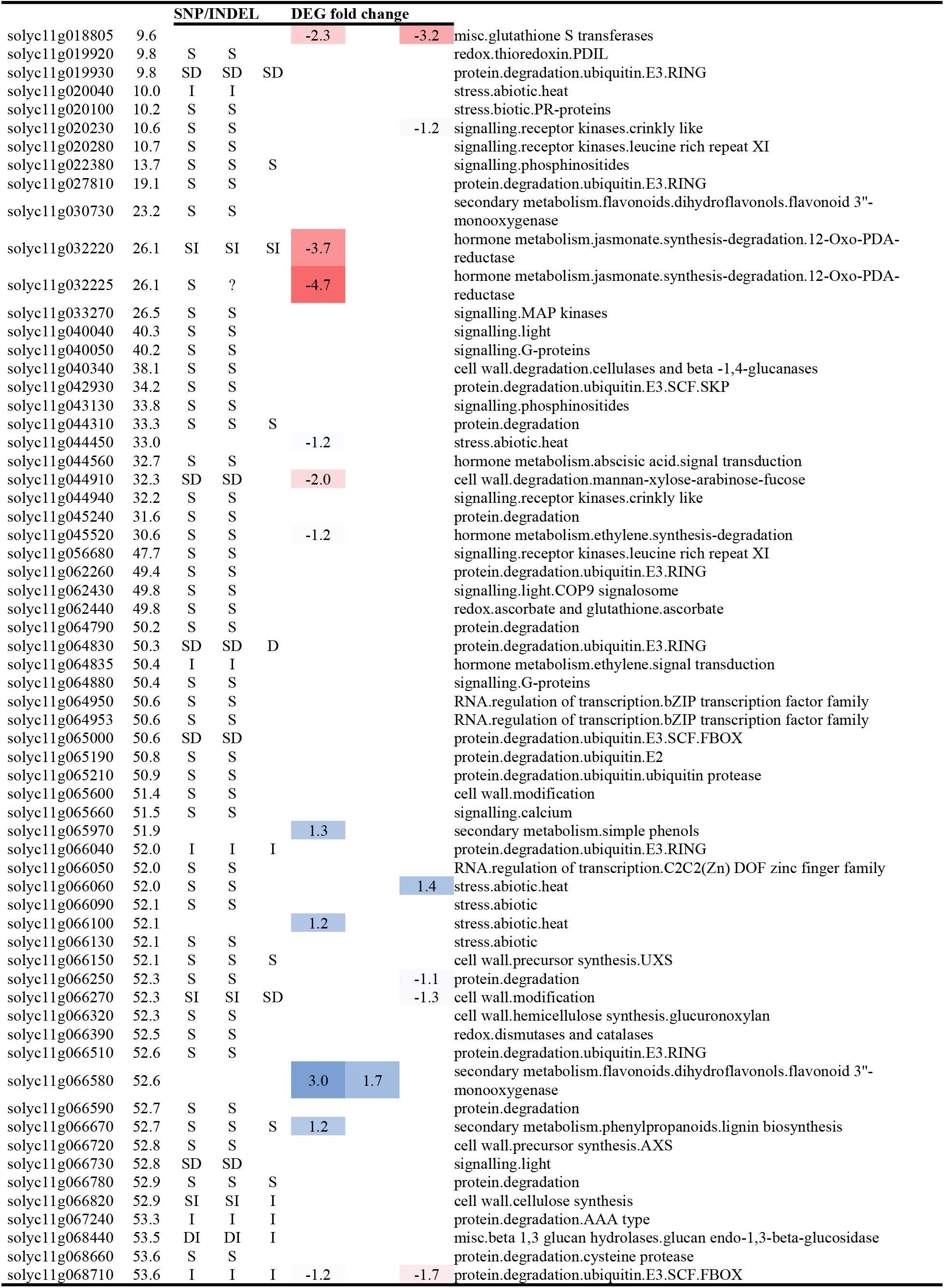
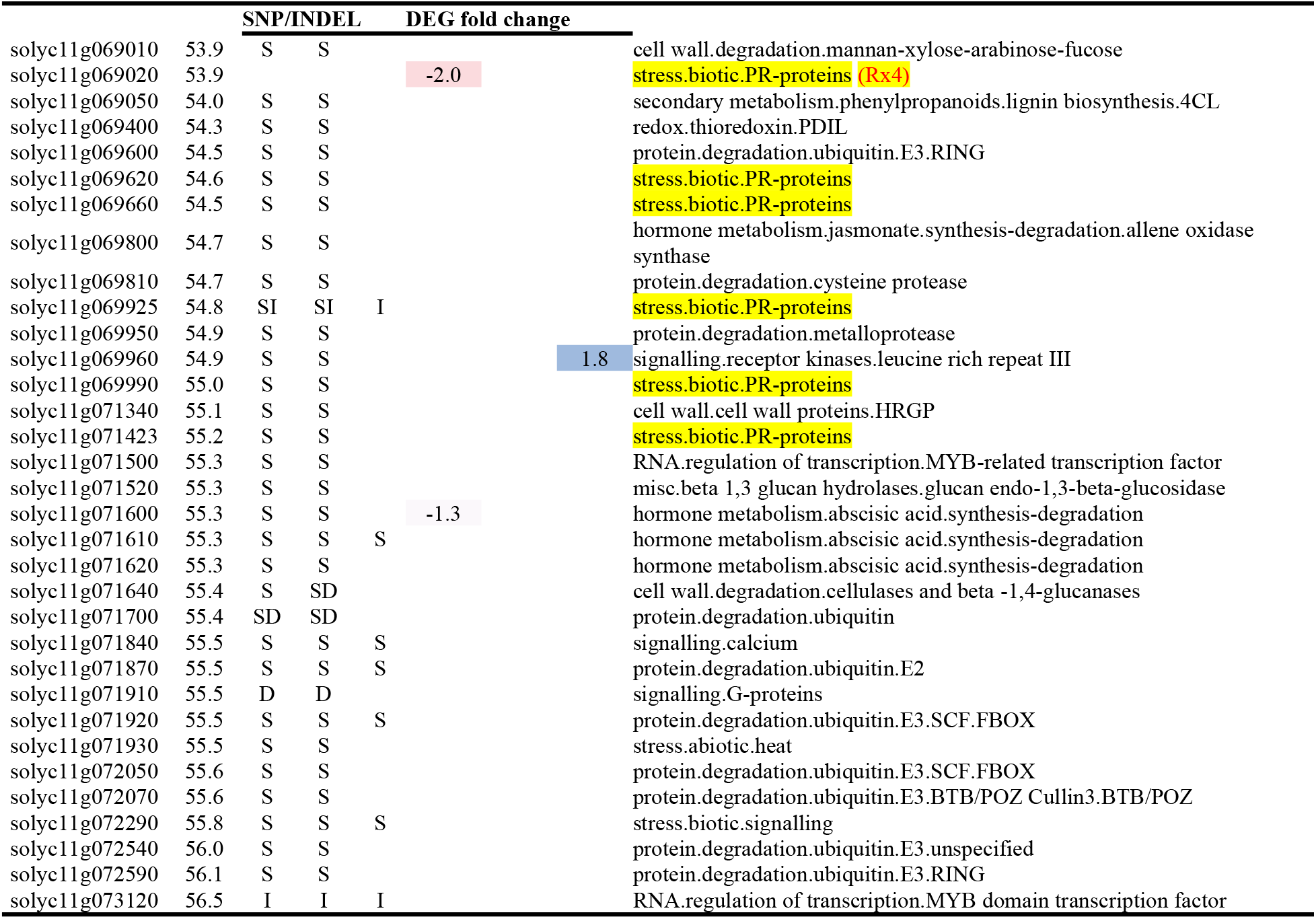
Biotic stress pathway DEGs or genes with SNP/INDELs on chr11. In the section of SNP/INDELs, “S,” “I” and “D” represent sequence variation in the homo form of SNP, INSERT and DELETION alternative to respective site in the reference genome of Heinz 1706. DEG fold changes are log_2_ of gene expression change induced by BS inoculation (IN/CK). Gene functions annotation bin were annotated by MapMan package in this study.

## Conclusions

We selected three distinct tomato breeding lines based on their different response to BS and performed RNA-Seq analysis for these lines to investigate differentially expressed genes (DEGs) induced two days after inoculation with BS race T4. Comparing functional involvements in various processes and past studies, we identified differentially expressed unique genes in resistance line PI 270443, such as those up-regulated PR-protein genes (solyc09g092300, solyc08g081790, and solyc09g092310) specific to this study. On another hand, a disease-associated R gene (solyc12g097000) was found down-regulated in susceptible NC 714. In addition to these differentially expressed genes, we have taken advantage of transcriptome-based RNA-Seq sequences to call SNP/INDELs from expressed genes between the three different tomato lines, and found that most of the molecular markers from resistant tomato lines PI 270443 and NC 1CELBR, were on chr11. Several PR-protein genes were identified in addition to known *Rx4* resistance genes. All these findings would be a valuable resource for tomato breeding aiming to develop resistant tomato to BS race T4.

## Acknowledgments

Help provided by Ann Piotrowski, Ragy Ibrahem, Ibrahim AlBallat, Mario De Jesus Velasco Alvarado, and Pragya Adhikari is highly appreciated. This research was partially supported by the College of Agriculture and Life Science (CALS) Agriculture Service of North Carolina State University, and National Science Foundation (grant# IOS-1546625) to Dilip R. Panthee.

## Supporting information

**Table S1. DEGs identified between inoculated and mock sample for each tomato lines**

**Table S2. Enriched GO terms**

**Table S3. SNP/INDELs across all chromosomes**

